# Environmental reservoir of resistance genes for the last resort antibiotic Cefiderocol

**DOI:** 10.1101/2025.05.26.656069

**Authors:** Remi Gschwind, Mehdi Bonnet, Anna Abramova, Victor Hugo Jarquín-Díaz, Marcus Wenne, Ulrike Löber, Nicolas Godron, Ioannis D. Kampouris, Faina Tskhay, Fouzia Naheed, Chloé Debroucker, Maximilien Bui-Hai, Inès El Aiba, Uli Klümper, Thomas U. Berendonk, Sofia K. Forslund-Startceva, Rabaab Zahra, Johan Bengtsson-Palme, Etienne Ruppe

## Abstract

Antibiotic resistance poses a global public health threat. Cefiderocol, a recently introduced siderophore cephalosporin, employs a “Trojan Horse” mechanism by exploiting bacterial iron uptake systems for cell entry. Yet, resistant clinical isolates are already observed in clinics and resistance mechanisms are difficult to characterize. Here, we applied functional metagenomics to identify cefiderocol resistance genes. Functional metagenomic DNA libraries from diverse environmental samples collected across several countries were expressed in a cefiderocol-sensitive *Escherichia coli* host. This yielded four resistant clones with DNA originating from wastewater or freshwater DNA libraries. The identified antibiotic resistance genes (ARGs) causing an increase in cefiderocol minimum inhibitory concentrations encoded for beta-lactamases (VEB-3, OXA-372 homolog and YbxI homolog) and a partial penicillin binding protein homolog. Three of four shared closest homologs in pathogenic bacteria. One ARG was associated with a mobile genetic element and was broadly distributed across all wastewater samples from every country surveyed. This study underscores the critical importance of environmental surveillance for ARGs, particularly for novel agents like cefiderocol with limited understanding of resistance mechanisms.

## Introduction

Antibiotic resistance represents a global public health threat, with resistant bacterial strains found in humans, animals and the environment, thereby reinforcing the ‘One Health’ concept ^1^. Beyond the dissemination of resistant strains across these compartments, bacteria can also transfer or acquire genetic material through horizontal gene transfer. The acquisition of antibiotic resistance genes (ARGs) can lead to the emergence of difficult-to-treat multidrug-resistant bacteria which were associated with 4.71 million deaths worldwide in 2021 ^2^. In 2024, the World Health Organization classified three extended-spectrum beta-lactamase (ESBL)-producing and seven carbapenemase-producing *Enterobacterales* among its 24 priority pathogens, due to their increasing resistance to last line antibiotics and widespread dissemination ^3,4^. In this context, the development of new therapeutic solutions is crucial. Cefiderocol, a recently developed antibiotic approved for clinical use, is a cephalosporin bearing a catechol group which acts as a siderophore by forming chelated complexes with ferric iron ^5^. Siderophores are molecules naturally produced by bacteria and secreted extracellularly to chelate iron. Employing a ’Trojan Horse’ strategy, iron-chelated cefiderocol utilizes the bacterial iron transport system to gain entry into the periplasm, where it disrupts cell wall biosynthesis by binding penicillin-binding proteins (PBPs).

Cefiderocol has demonstrated efficacy against a broad spectrum of Gram-negative bacterial isolates, including ESBL and carbapenemase producers ^6–9^. Its stability against hydrolysis by class D beta-lactamases (OXA-48, OXA-40, OXA-23) and other class A and class B beta-lactamases (IMP-1, VIM-2, NDM-1, KPC-2, KPC-3, L1) has been established ^10,11^. However, cefiderocol-resistant isolates have been detected, even in patients without prior exposure to the antibiotic ^12^. Cloning and expression studies have shown that certain beta-lactamases, including class A (KPC, PER, SHV, BEL), class B (NDM), class C (AmpC) and class D (OXA) enzymes, are associated with decreased cefiderocol susceptibility ^13–19^. This can be attributed to the ability of these beta-lactamases to hydrolyze or trap cefiderocol, as observed with some KPC variants ^20^. Other resistance mechanisms involve target modifications. For instance, YRIN or YRIK insertions at position 338 in the PBP3 encoding gene have been linked to elevated cefiderocol minimum inhibitory concentrations (MIC; 8, 21, 22). Membrane permeability modifications can also influence cefiderocol MICs ^15^. Furthermore, mutations or deletions in genes involved in iron uptake can impact the cefiderocol resistance phenotype ^8,9,12,21,23–29^. Notably, cefiderocol resistance is associated not only with gene mutations or truncations but also with variations in gene expression ^30^. This multifactorial nature of cefiderocol resistance complicates its study, making cloning and expression experiments the gold standard for elucidating the contribution of individual mechanisms.

Efforts are increasingly directed towards improving our understanding and monitoring strategies of ARGs ^31^. Metagenomics facilitates the detection of ARGs within an environment or a bacterial strain by comparing sequences to ARG databases. However, data concerning the effects of ARGs on novel antibiotics, such as cefiderocol, is often limited. Functional metagenomics can address this gap by identifying ARGs not yet described in existing databases or by providing phenotypic data for known ARGs ^32^. This technique identifies ARGs based on phenotype rather than solely on sequence homology. It is performed by expressing DNA libraries in antibiotic-sensitive hosts in the presence of antibiotics. Resistant clones harbor a DNA fragment containing an ARG. Its sequence can be determined to increment ARG databases like ResFinderFG, a database of ARG identified by functional metagenomics ^33^. Furthermore, the resistant clones can be used to precisely characterize the associated resistance phenotype across an entire antibiotic family.

Although cefiderocol resistance has been primarily studied in clinical strains, data regarding its prevalence in the environment remains scarce. Nonetheless, the environment is a crucial compartment within the ‘One Health’ framework. While bacteria and their ARGs are transferred between compartments, the environment, characterized by its extensive niche diversity, harbors a highly diverse reservoir of genes including ARGs ^34^. Environmental bacteria are considered the ancestral hosts for most ARGs, predating clinical antibiotic use, and the environment still serves as a shared source of ARGs to both environmental and pathogenic strains ^35,36^. In this study, we aimed to employ functional metagenomics to identify ARGs conferring cefiderocol resistance across diverse environmental samples.

## Results

A total of 47 samples were collected for the EMBARK project (**Table 1**), 17 from Sweden, two from France, 16 from Germany and 12 from Pakistan. DNA from each sample was extracted and either subjected to metagenomic sequencing only or metagenomic sequencing and functional metagenomics depending on DNA concentrations and due to DNA requirements of functional metagenomics (at least 800 ng). The whole study design can be found in **Figure 1**.

**Figure 1:**
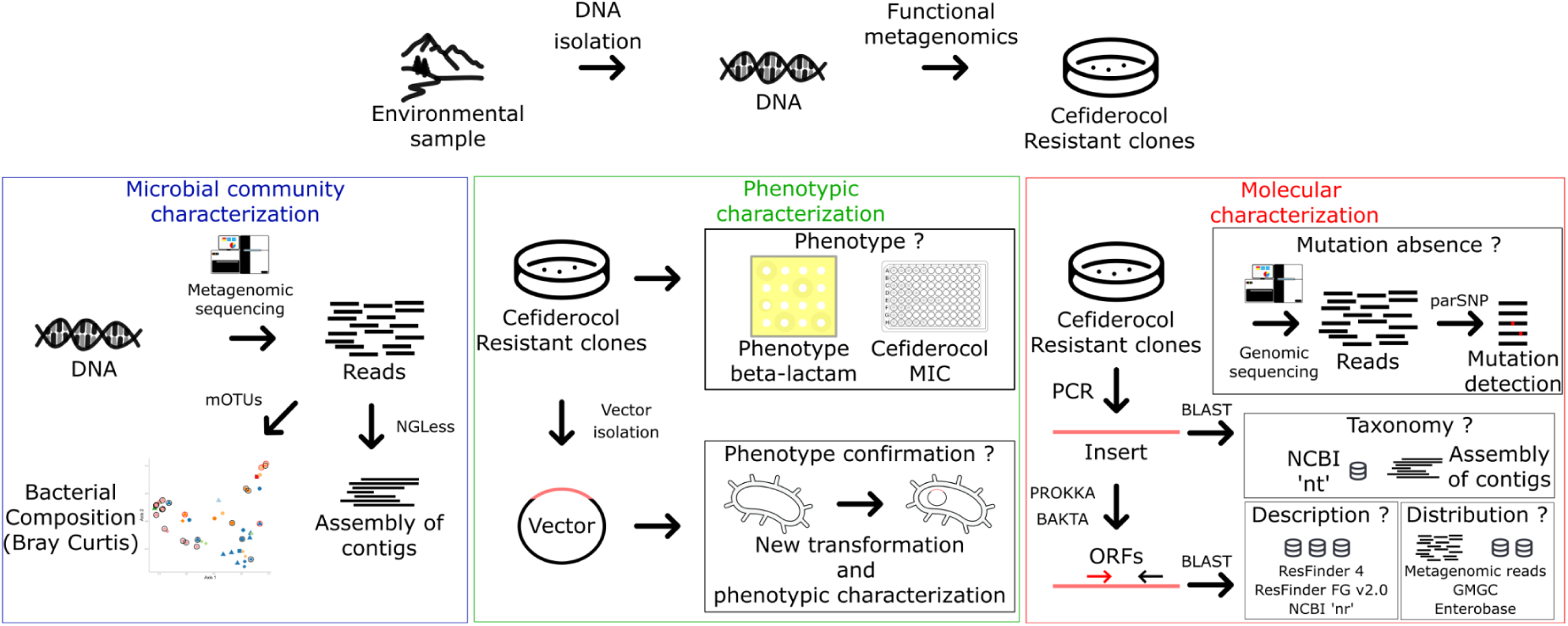
Study design.

**Table 1:**
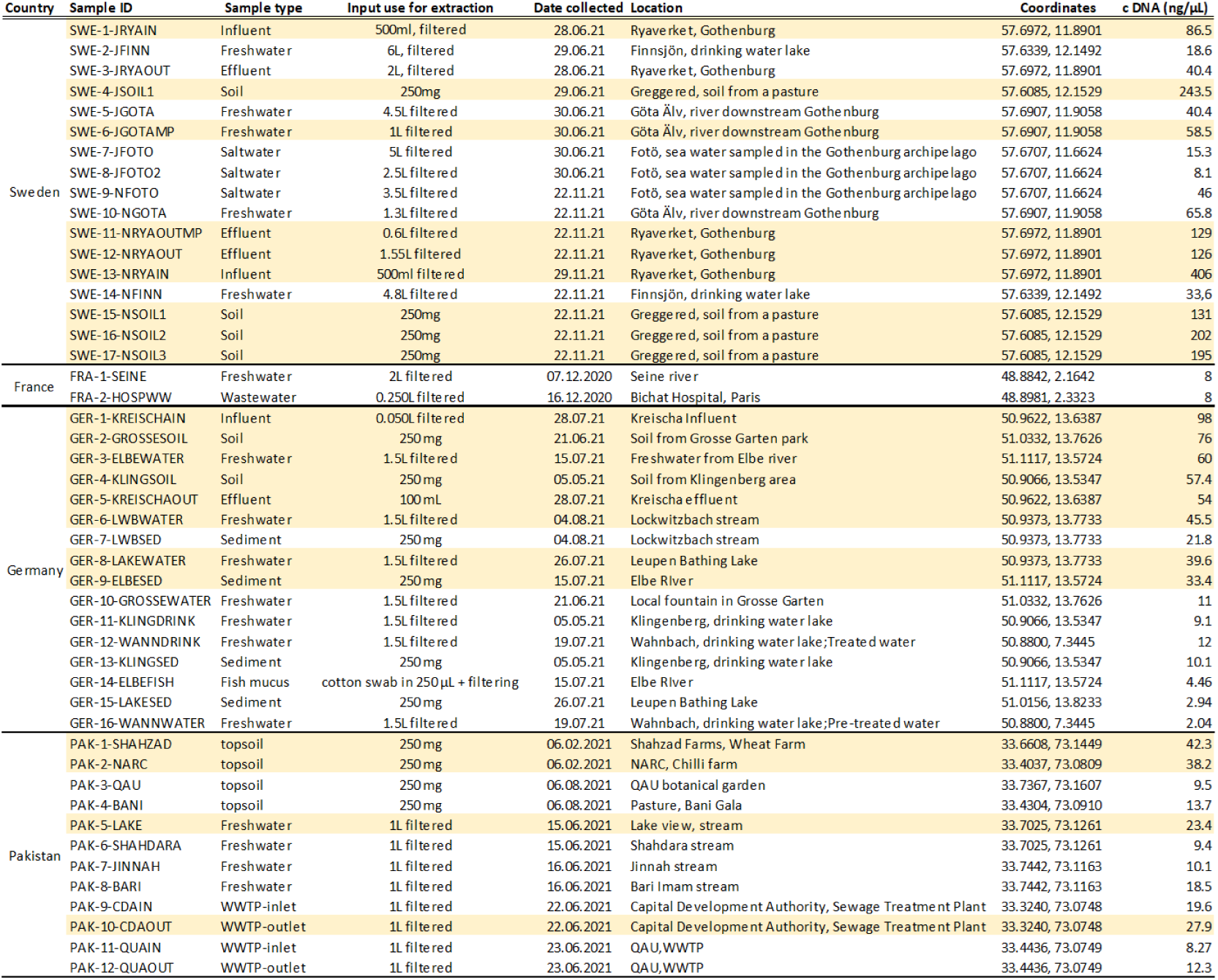
Samples included in the EMBARK project. Samples which were included in the functional metagenomics analysis are highlighted in yellow.

### Functional metagenomics libraries production and selection of cefiderocol resistant clones

Due to substantial requirements regarding DNA quantity, 21 samples were selected for functional metagenomics analysis of cefiderocol resistance. DNA libraries were constructed via a tagmentase shearing process and Gibson cloning in a pHSG299 expression vector. Each library was then expressed in a cefiderocol-sensitive K12 *Escherichia coli* followed by selection of cefiderocol-resistant clones on LB agar media containing cefiderocol (1 mg/L). Resistant clones were detected in four samples (19%): three wastewater samples (SWE-1-JRYAIN, GER-1-KREISCHAIN and GER-5-KREISCHAOUT) and one freshwater sample (GER-3-ELBEWATER).

### Shotgun metagenomics analysis of EMBARK samples

To evaluate if the identification of DNA fragments responsible for cefiderocol resistance might be correlated with specific bacterial communities, all DNA extracts were subjected to shotgun metagenomic sequencing. To this end, taxonomic profiling of the metagenomes was done using mOTU v3.1 ^37^ and a PCoA analysis of beta-diversity using Bray-Curtis dissimilarity was performed (**Figure 2**). This analysis effectively differentiated samples based on their environmental origin (wastewater input or output, soil, and freshwater). Samples that tested positive for cefiderocol resistance via functional metagenomics did not exhibit clustering within any specific environmental category based on their bacterial community composition.

**Figure 2:**
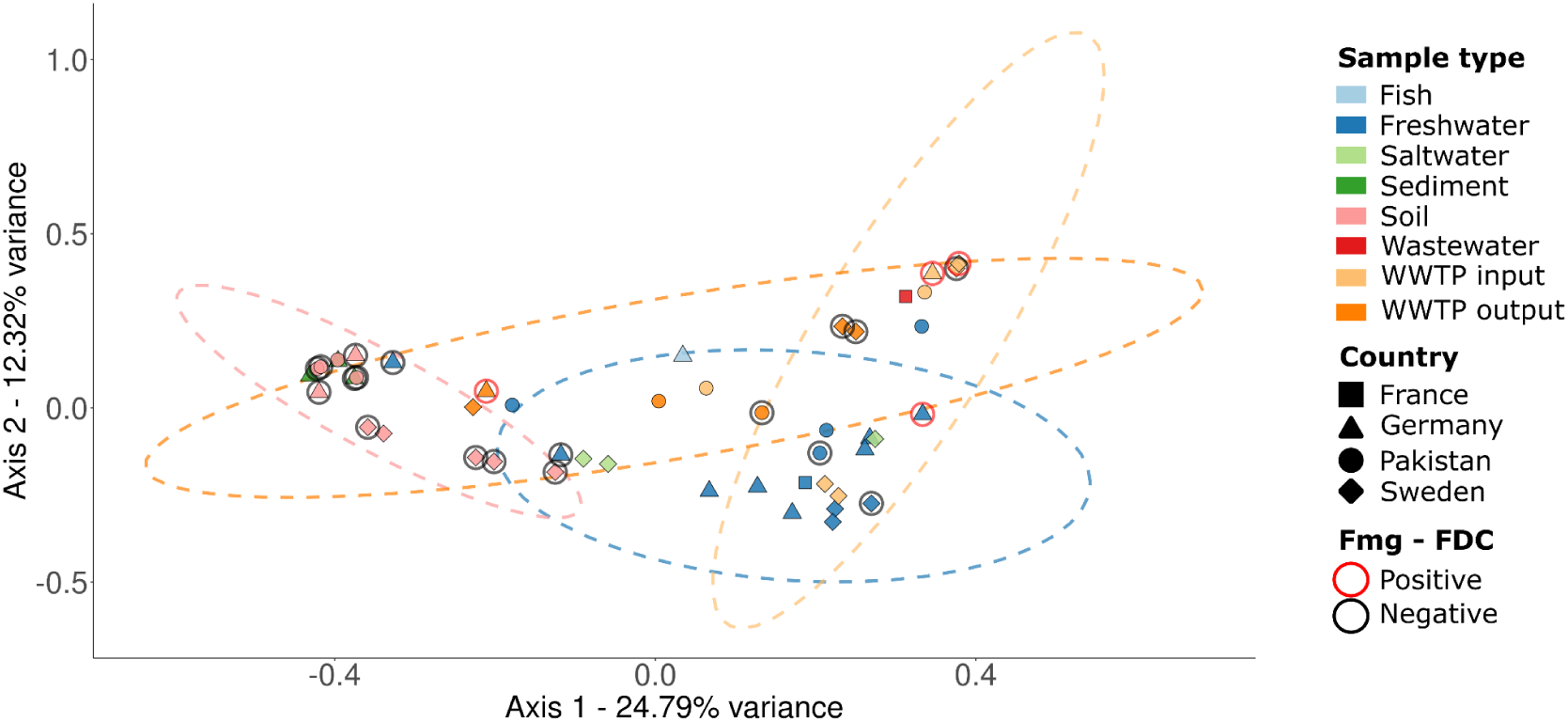
PCoA analysis of the bacterial composition of each sequenced sample based on Bray-Curtis dissimilarity. Samples are classified based on their type, country of origin, whether they were analyzed with functional metagenomics and whether they were positive or negative for cefiderocol resistance. Data ellipses were computed for sample types with enough data points (i.e. freshwater, soil, WWTP input and WWTP output) using the function stat_ellipse() from ggplot2, with default statistical parameters (assuming multivariate t-distributions). WWTP: wastewater treatment plant; Fmg-FDC: cefiderocol resistance studied using functional metagenomics.

### Cefiderocol-resistant clones characterization

The cefiderocol-resistant phenotype of each clone was confirmed. First, this was achieved by comparing the phenotypic data with fresh competent K12 *E. coli* transformed with the same plasmid containing the ARG extracted from the resistant clone. Second, by sequencing the genomic DNA of each resistant clone to confirm the absence of mutations that could contribute to an increased cefiderocol MIC (see **Supplementary Data**). The observed cefiderocol resistance was indeed due solely to the expression of the DNA fragment cloned into the expression vector for the four resistant clones. Each ARG containing DNA fragment cloned into the expression vector and responsible for cefiderocol resistance was amplified by PCR and Sanger sequenced. It was characterized molecularly using BLASTN or BLASTP with several databases (‘nt’, ‘nr’, ResFinderFG v2.0, ResFinder v4.6.0; 33, 38) and open reading frames (ORFs) were predicted using PROKKA v1.14 and BAKTA v1.11 ^39,40^. Resistant clones were also characterized phenotypically using cefiderocol MIC determination and disc diffusion assay.

A cefiderocol-resistant clone was found using functional metagenomic DNA library from an influent wastewater sample from Ryaverket, Gothenburg, Sweden (ID: SWE-1-JRYAIN). The DNA fragment responsible for cefiderocol resistance had a size of 1,316 bp (**Table 2, Supp Figure 2**). The closest homolog in the ‘nt’ database was a sequence found in *Citrobacter freundii* (91.4% identity and 71% coverage) followed by other sequences from Gammaproteobacteria such as *Morganella morganii* or *Pseudomonas indoloxydans* (97,2% identity and 59% coverage). Two ORFs were detected. One encoded a class D beta-lactamase annotated as OXA-10 beta-lactamase or OXA-372 family carbapenem-hydrolyzing class D beta-lactamase (by PROKKA or BAKTA, respectively). The other ORF was annotated as a hypothetical protein by both programs. The class D beta-lactamase was presumed to be responsible for the cefiderocol resistance phenotype. A homolog (*bla*_OXA-372_, 93.2% identity and 100% coverage) was found in the ResFinder 4.6.0 database. Homologous proteins expressed by *Pseudomonas aeruginosa*, *M. morganii* and *Ectopseudomonas oleovorans* were identified in the NCBI ‘nr’ database (identity >96%, coverage >99%). Phenotypically, the clone exhibited a cefiderocol MIC of 2 mg/L, a 16-fold increase compared to the K12 strain transformed with an empty expression vector. This clone also resisted most of the beta-lactam antibiotics tested, except for cefalotin, cefoxitin and carbapenems (**Figure 3A**). Combinations of beta-lactam antibiotics and beta-lactamase inhibitors, including ceftazidime-avibactam and ceftolozane-tazobactam, were ineffective.

**Figure 3:**
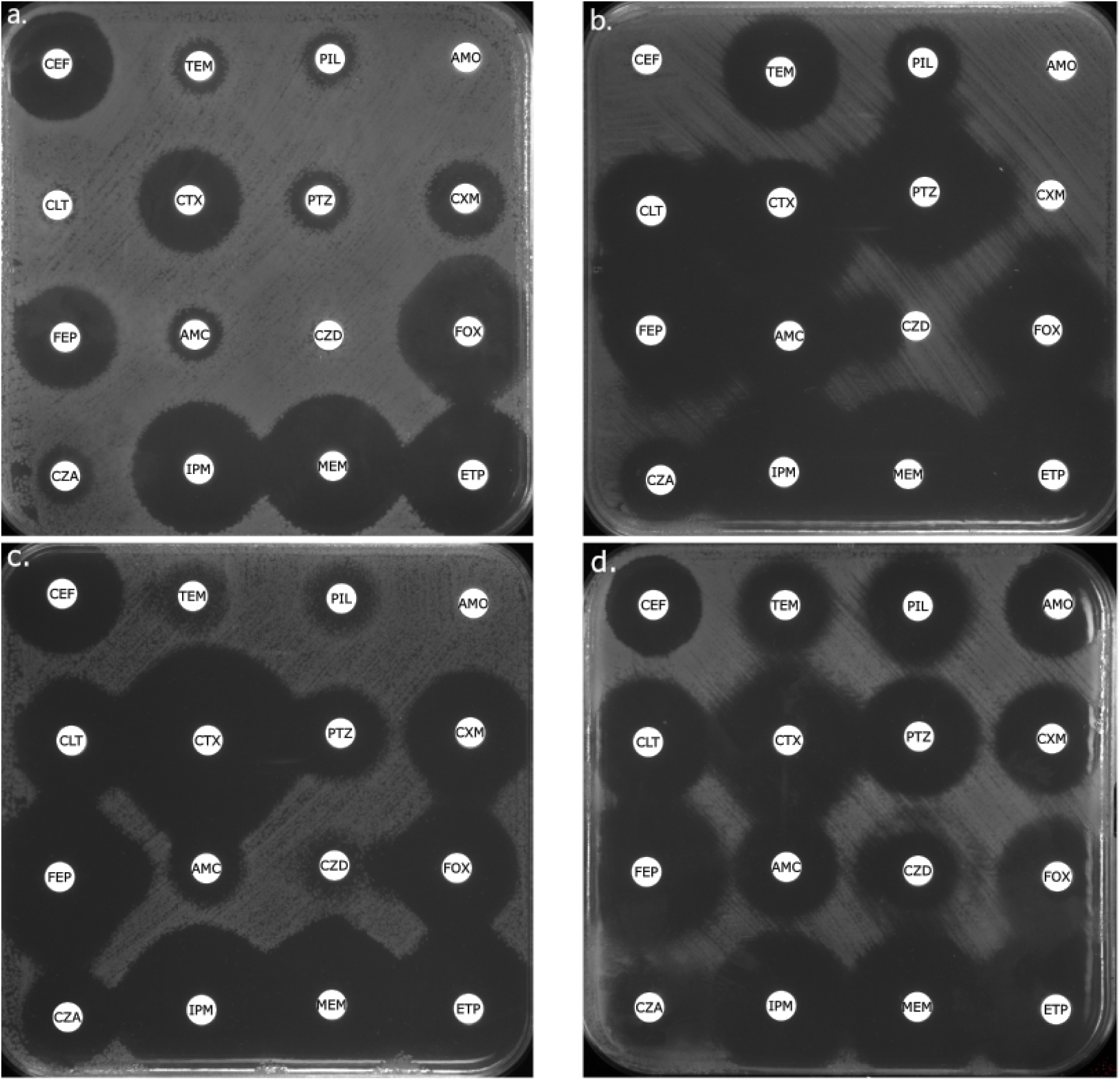
Disc diffusion assay for each cefiderocol-resistant clones. **A** SWE-1-JRYAIN; **B** GER-1-KREISCHAIN; **C** GER-3-ELBEWATER; **D** GER-5-KREISCHAOUT; CEF: cephalotin (30 µg); TEM: temocillin (30 µg); PIL: piperacillin (30 µg); AMO: amoxicillin (20 µg); CLT: ceftolozane (30 µg) + tazobactam (10 µg); CTX: cefotaxime (5 µg); PTZ: piperacillin (30 µg) + tazobactam (6 µg); CXM: cefuroxime (30 µg); FEP: cefepime (30 µg); AMC: amoxicillin (20 µg) + clavulanic acid (10 µg); CZD: ceftazidime (10 µg); FOX: cefoxitine (30 µg); CZA: ceftazidime (10 µg) + avibactam (4 µg); IPM: imipenem (10 µg); MEM: meropenem (10 µg); ETP: ertapenem (10 µg).

**Table 2:**
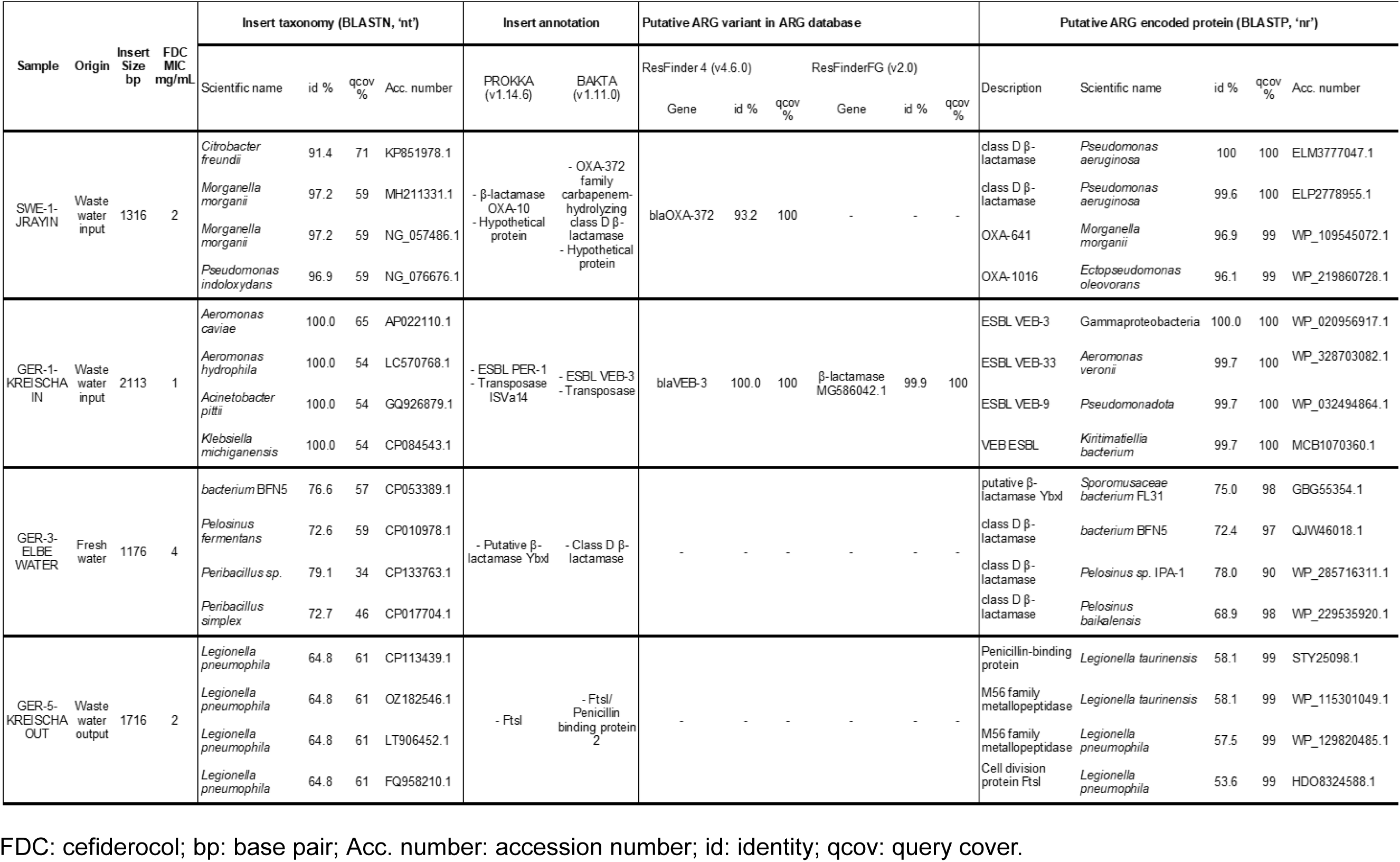
Molecular characteristics of insert sequences and its ARGs responsible for cefiderocol MIC increase.

A second cefiderocol-resistant clone was obtained using functional metagenomic DNA library from another influent wastewater sample from Kreischa, Germany (ID: GER-1-KREISCHAIN). The DNA fragment causing cefiderocol resistance was a 2,113 bp insert (**Table 2, Supp Figure 2**). Its closest homologs (identity percentage 100%) in ‘nt’ database were sequences found in Gammaproteobacteria (*Aeromonas caviae*, *Aeromonas hydrophila*, *Acinetobacter pittii* and *Klebsiella michiganensis*), albeit with a partial coverage (ranging from 52 to 63%). The covered region was characterized by an ORF presumed to confer cefiderocol resistance, annotated as an ESBL PER-1 by PROKKA or as an extended-spectrum class A beta-lactamase VEB-3 encoding gene by BAKTA. The remaining sequence of the insert was annotated as an IS4 family transposase, potentially explaining the partial coverage observed with homologous sequences in the NCBI ‘nt’ database. In ResFinder 4.6.0, the VEB-3 class A ESBL annotation was confirmed, with the gene mapping perfectly to *bla*_VEB-3_, (100.0% identity and 100% coverage). Homologs were also identified in the ResFinderFG v2.0 database (a beta-lactamase identified in antibiotic-polluted stream sediment, 98.0% identity and 100% coverage) and the NCBI ‘nr’ database (VEB beta-lactamase from Gammaproteobacteria, 100% identity, 100% coverage). The cefiderocol-resistant clone displayed an ESBL profile with synergistic effects observed for inhibitors such as clavulanic acid or tazobactam and cefotaxime, ceftazidime and cefepime (**Figure 3B**). Cefoxitin, temocillin and carbapenems remained effective. Notably, a synergistic effect was observed between cefoxitin and cefuroxime, as well as ceftazidime. Its MIC against cefiderocol was 1 mg/L, an 8-fold increase.

The third cefiderocol-resistant clone was identified using a functional metagenomic DNA library originating from a freshwater sample of the Elbe River downstream of Dresden, Germany (ID: GER-3-ELBEWATER). The DNA fragment causing cefiderocol resistance was 1,176 bp long (**Table 2, Supp Figure 2**). In the NCBI ‘nt’ database, the closest homologs were an unclassified bacterium (*bacterium* BFN5, 76.6% identity 57% coverage) and bacteria from the Bacillota phylum (*Peribacillus* genera, and *Pelosinus fermentans*; 72.6-79.1% identity, 34-59% coverage). Annotation tools identified a single ORF annotated as a class D beta-lactamase or a putative beta-lactamase YbxI encoding gene. This ARG was not found in any ARG database tested. The closest protein homologs in the NCBI ‘nr’ database were class D beta-lactamases from Bacillota with 68.9 to 78.0% identity, although with a high coverage (ranging from 90 to 98%). The GER-3-ELBEWATER resistant clone exhibited resistance to penicillin, ceftazidime and temocillin. A synergistic effect between cefoxitin and ceftazidime was also observed (**Figure 3C**). Clavulanic acid and avibactam partially restored the activity against amoxicillin and ceftazidime, respectively. The cefiderocol MIC was 4 mg/L, a 32-fold increase.

The final cefiderocol-resistant clone was obtained using a functional metagenomic DNA library from a fresh water sample (ID: GER-5-KREISCHAOUT) collected downstream of the same wastewater treatment plant as GER-1-KREISCHAIN. Its insert sequence, 1,716 bp in length, exhibited 64.8% identity (61% coverage) with a *Legionella pneumophila* sequence (**Table 2, Supp Figure 2**). Annotation tools identified a single ORF as peptidoglycan D,D-transpeptidase FtsI encoding gene or as cell division protein FtsI/PBP2 encoding gene. No homologous sequences were found in tested ARG databases. The closest protein match in the ‘nr’ database was a PBP found in *L. taurinensis* (58.1% identity and 99% coverage). Phenotypically, this clone displayed an atypical profile, exhibiting low-level resistance only to ceftazidime, with synergistic effect observed between ceftazidime and clavulanic acid or cefoxitin (**Figure 3D**). A synergistic effect was observed between imipenem and ceftazidime-avibactam. Its cefiderocol MIC was 2 mg/L, a 16-fold increase. Given the unclear resistance mechanism, further investigations were performed. To check for potential beta-lactamase-mediated resistance, a nitrocefin hydrolysis test and cefiderocol-avibactam MIC were performed. The nitrocefin hydrolysis test was negative and cefiderocol MICs did not differ with or without avibactam. The 3D structures of several proteins were predicted using AlphaFold (41): FtsI from *E. coli* (accession number: WP_000625659.1) with or without YRIN/YRIK insertions at position 338, the protein identified by functional metagenomics, and various PBPs found in *L. pneumophila* (accession number: CP013742). A conserved domain within each FtsI or PBP2-encoding gene was aligned using PyMol to the protein identified by functional metagenomics (**Figure 4A, B, C**; 42). Yet, it showed that the protein encoded by GER5 might lack a domain compared to other PBP or FtsI encoding genes. We therefore hypothesised that the gene might have been truncated at the shearing step of the functional metagenomic library preparation process. To test this hypothesis, contigs were assembled from the metagenomic data of GER-5-KREISCHAOUT sample using NGLess ^43^ and MEGAHIT ^44^ and the FtsI encoding gene identified was aligned to the assembled contigs. This gene was found on a 23,400 bp contig with 882 additional base pairs due to a start codon found upstream of the break induced by the shearing step during functional metagenomic library preparation (see **Supp Figure 3**). Protein structure alignments revealed the highest similarity between the short and extended version of the protein identified through functional metagenomics (**Figure 4D**). The second-best alignment was with *E. coli* FtsI, followed by *L. pneumophila* PBP2. Phenotypic analysis, including cefiderocol MIC determination and disc diffusion assays, showed no difference between *E. coli* K12 carrying the extended gene version and the one carrying the shorter version.

**Figure 4:**
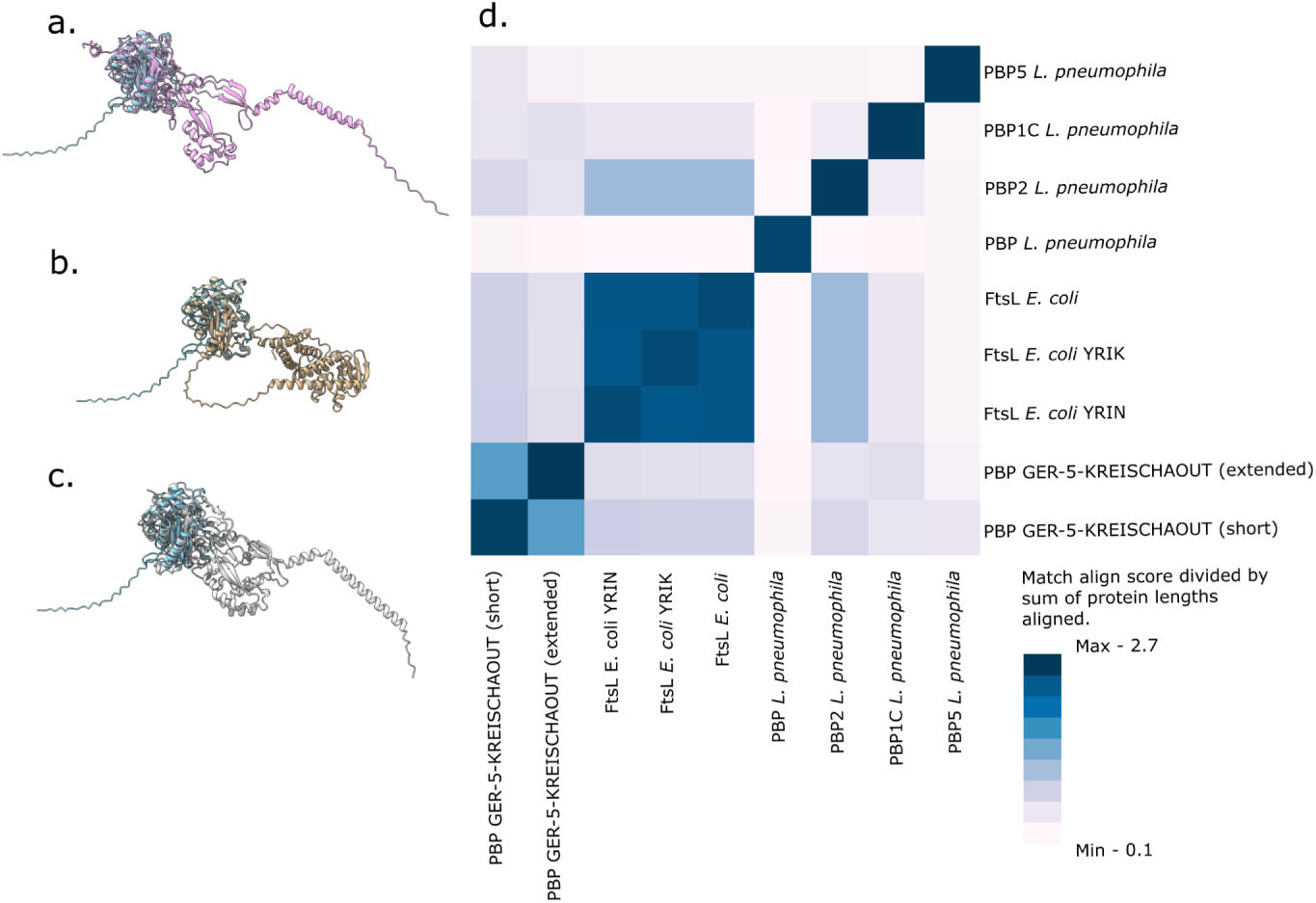
3D structures predicted by AlphaFold 3 and aligned using PyMOL. **A** FtsI from GER-5-KREISCHAOUT (short) *vs E. coli* FtsI; **B** FtsI from GER-5-KREISCHAOUT (short) *vs* FtsI from GER-5-KREISCHAOUT (extended); **C** FtsI from GER-5-KREISCHAOUT (short) *vs L. pneumophila* PBP2; **D** Match align score divided by sum of protein lengths aligned.

### Cefiderocol ARGs distribution

The distribution of cefiderocol resistance genes identified through functional metagenomics was investigated in sample metagenomic reads (**Figure 5**), in GMGC and in EnteroBase ^45,46^. With regards to geographical distribution, *bla*_VEB-3_ from GER-1-KREISCHAIN was detected in all countries and was the most prevalent cefiderocol resistance gene (16/47 positive samples), with relative abundance up to 1.5E-5 (mapped reads/total reads) in samples from Pakistan. This was followed by the oxacillinase-encoding gene from SWE-1-JRYAIN, which was also found in every country but at lower abundances than *bla*_VEB-3_. These two genes exhibited a broad distribution in wastewater and freshwater samples (18/36), with all wastewater samples testing positive for at least one cefiderocol resistance gene. The PBP-encoding gene found in the GER-5-KREISCHAOUT wastewater sample was also present in freshwater (GER-6-LWBWATER) but showed a local distribution in the Dresden region of Germany. The cefiderocol-resistance gene identified in GER-3-ELBEWATER was not detected in any sample. None of the cefiderocol-resistant genes identified through functional metagenomics was found in soil sample sequencing reads. When we checked the environmental distribution within the GMGC, the cefiderocol resistance genes identified in SWE-1-JRYAIN, GER-3-ELBEWATER and GER-5-KREISCHAOUT, encoded proteins with low identity percentage (%id ranging between 53.9 and 68.9%) to their closest homologs in GMGC. The closest homolog to the GER-1-KREISCHAIN gene was the GMGC10.001_990_215.UNKNOWN unigene from *P. aeruginosa*. This unigene was found in 2/7,059 human gut samples, 2/1,139 human skin samples and 3/22 wastewater samples. Regarding the host distribution in EnteroBase, only homologs of *bla*_VEB-3_ from GER-1-KREISCHAIN were identified in nine *E. coli* samples (97.8 to 97.9% identity and 100% coverage; **Supp Table 1**). These *E. coli* isolates belonged to phylogroup C (three isolates) and A (six isolates), and included sequence types ST176 (four isolates), ST88 (one isolate), ST472 (one isolate), ST10 (one isolate), ST471 (one isolate) and ST410 (one isolate).

**Figure 5:**
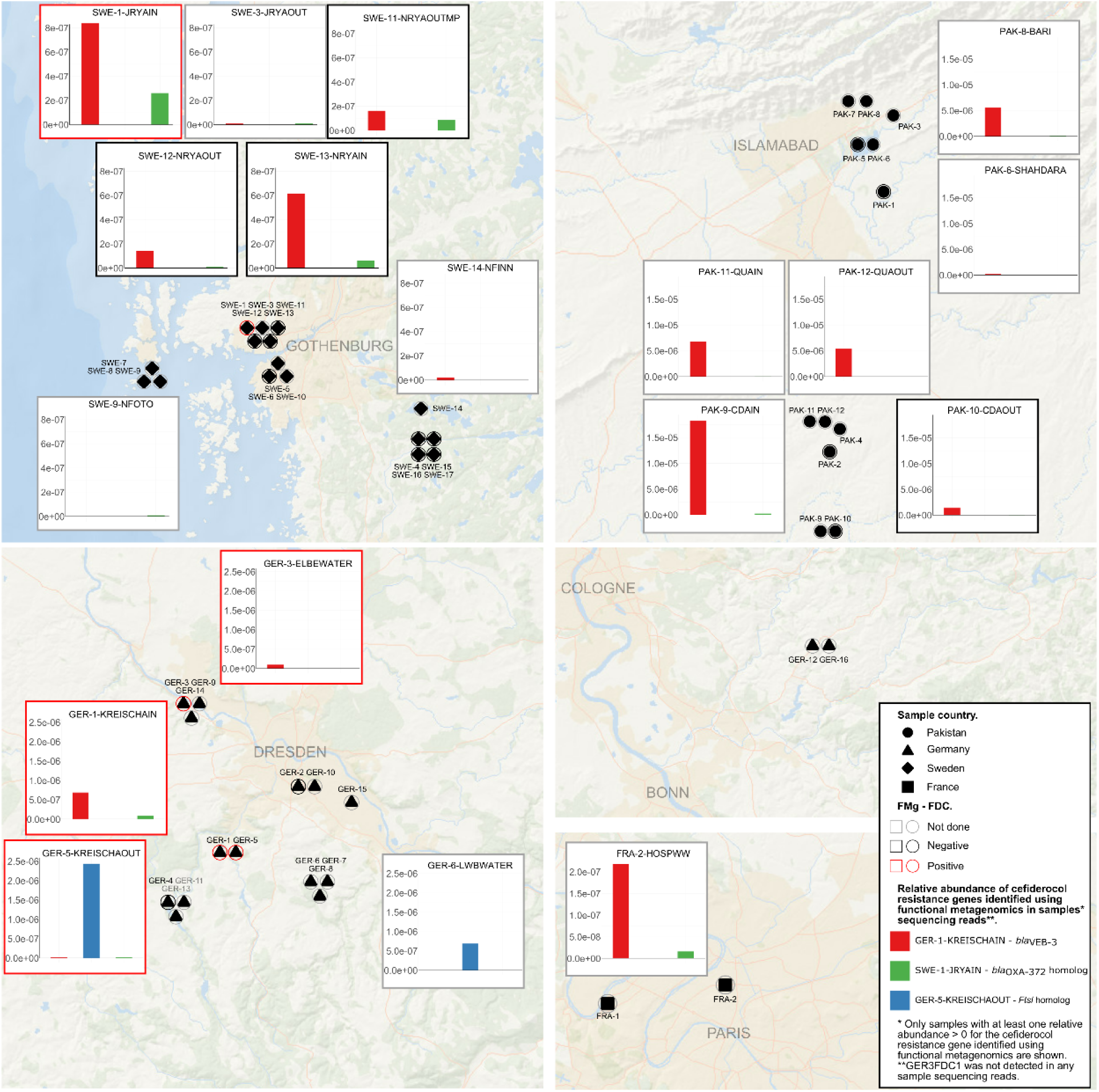
Map showing the geographic locations of samples included in functional metagenomics studies of cefiderocol resistance and the relative abundance of the genes identified using functional metagenomics in whole metagenome reads (if at least one had a relative abundance >0 in sequencing reads).

## Discussion

Using functional metagenomics, our study established the presence of four distinct ARGs conferring resistance to cefiderocol, a recently-released, last line antibiotic, within environmental samples, most likely devoid of any selection-inducing cefiderocol exposure.

Three of the identified genes encoded beta-lactamases or beta-lactamases homologs while one encoded a partial PBP homolog. The VEB-3 beta-lactamase conferred an 8-fold increase in cefiderocol MIC. Previously, VEB beta-lactamases were not associated with cefiderocol resistance. For instance, the VEB-1 beta-lactamase was not found to hydrolyze cefiderocol in enzymatic assays nor increase cefiderocol MIC, either in clinical isolates or when cloned into sensitive *E. coli* ^8,18^. *P. aeruginosa* clinical isolates producing VEB beta-lactamases exhibiting cefiderocol MIC >1mg/L were reported, yet additional resistance mechanisms may have contributed to the observed MIC increase ^9^. For Enterobacterales isolates, for example, cefiderocol at 2 mg/L inhibited around 80% of isolates. The most significant drop was seen in isolates producing NDM beta-lactamase (41% inhibited isolates) and in isolates with combinations of ESBL production and porin loss ^6^. Thus, to our knowledge, our findings are the first to support that a VEB beta-lactamase can increase the cefiderocol MIC by itself. We also identified two class D beta-lactamases (an OXA-372 and an YbxI variant) responsible for a 16 to 32-fold increase in cefiderocol MIC. Enzymatic assays indicated that OXA-48, OXA-40 and OXA-23 beta-lactamases do not directly hydrolyze cefiderocol, their expression in *E. coli* and in *Acinetobacter baumannii* (including OXA-58 in the latter) also failed to elicit an increased cefiderocol MIC ^10,11,18^. Only OXA-427 conferred an 8-fold MIC increase when cloned in *P. aeruginosa* and this gene has also been associated with cefiderocol resistance in clinical isolates ^47^. From environmental samples, while other mechanisms might be involved, OXA-181 identified by PCR in *Enterobacter cloacae* complex was also associated with increased cefiderocol MIC ^48^. The YbxI beta-lactamase was described in *Bacillus subtilis* as a low activity beta-lactamase and had not been previously associated with cefiderocol resistance nor elevated cefiderocol MICs ^49^.

Besides beta-lactamases, we identified a partial PBP-encoding gene responsible for a 16-fold increase in cefiderocol MIC. Expression of this truncated gene was sufficient to cause a MIC increase. Protein structural analysis revealed the absence of a specific domain. Yet, the complete version of the protein was also expressed in K12 *E. coli* and showed the same phenotypic profile. The 3D structure of either the partial or complete protein exhibited the highest similarity to *E. coli* PBP-3, a primary target of cefiderocol ^8^. PBP-3 harboring YRIN or YRIK insertion in position 338 have been linked to cefiderocol resistance ^8,12,21,22^. However, these insertions were absent in both the partial and complete version of the gene identified in our clone or metagenomic contig and the protein did not exhibit better alignment to PBP-3 containing YRIN or YRIK insertions.

Of the 21 environmental samples analyzed using functional metagenomics, four (19%) yielded cefiderocol resistance genes. Using whole metagenomic sequencing though, these genes were detected in 18 out of 47 environmental samples (38%). Some genes exhibited a broad distribution (*bla*_VEB-3_, likely due to associations with mobile genetic elements), while others showed localized distribution (PBP encoding gene from GER-5-KREISCHAOUT) or were not detected (class D beta-lactamase encoding gene from GER-3-ELBEWATER). Notably, the cefiderocol resistance gene from GER-1-KREISCHAIN was found in 100% of wastewater metagenomic sequencing reads. Wastewater, known to harbor diverse microbial communities including pathogenic bacteria and their genetic content (ARGs, virulence factors, mobile genetic elements), has been identified as a hotspot for mobilization and promotion of ARGs ^34,50^. In contrast, while soil is often considered a potential reservoir for resistance ^36^, soil samples exhibited no detectable cefiderocol resistance genes, neither through functional metagenomics nor by analyzing the distribution of cefiderocol resistance genes identified in other samples using functional metagenomics. The origins of antibiotic resistance in soil are frequently linked to naturally occurring antibiotics produced by soil bacteria and fungi. Cefiderocol, as a novel, synthetic antibiotic, likely has a limited environmental presence in soil or environments with limited anthropogenic contact. Moreover, its use as a last resort antibiotic, suggests a minimal environmental exposure minimizing the potential selection of specific cefiderocol resistance genes. Here, the identified ARGs conferred resistance not only to cefiderocol but also to other beta-lactam antibiotics more commonly used in human medicine. Therefore, broader resistance profiles could explain selection of cefiderocol resistant genes and prevalence in environments like wastewater, which receive diverse human-derived antibiotics. It is noteworthy that the genes identified using functional metagenomics had homologs or were found in bacterial isolates associated with clinical settings. Specifically, the *bla*_VEB-3_ gene can be identified in wastewater samples, in freshwater samples and in clinical context ^51,52^. Homologs were identified in genomes from the Enterobase database, notably in ST410 *E. coli* strains, a disseminating ST, considered a high-risk multi-drug resistant clone causing human disease ^52–56^. Furthermore, the closest homolog of the oxacillinase-encoding gene from SWE-1-JRYAIN was found in *C. freundii*, a bacterium known for its pathogenic potential ^57^.

Functional metagenomics, while powerful, possesses inherent limitations. It does not provide an exhaustive description of all ARGs within a sample. This is exemplified in our study, where cefiderocol resistance genes identified through functional metagenomics were not consistently detected in corresponding metagenomic sequence data, and genes detected in metagenomic sequence data were not always found by functional metagenomics. Nevertheless, functional metagenomics remains crucial for elucidating resistance mechanisms to novel antibiotics. Phenotypic cefiderocol resistance in bacterial isolates has been documented, but its mechanistic basis is often complex, involving multiple ARGs, virulence factors, and/or mutations. Traditional approaches, such as cloning individual ARGs into a susceptible host, are exceedingly laborious. Such approaches involve the sequential testing of each gene, a daunting task given the hundreds of potentially involved ARGs, and often result in negative outcomes, (no increase in MIC). Conversely, functional metagenomics eliminates the need for prior assumptions. It allows for the cloning of large DNA fragments without prior knowledge of their function or identity, and directly selects genes conferring increased MICs. This approach potentially accelerates the identification of new phenotypic information regarding known ARGs and the discovery of novel resistance genes. Moreover, as evidenced here, it could also identify more precisely specific gene regions or encoded protein domains necessary for the acquisition of the resistance phenotype. Our findings revealed ARGs not previously linked to this resistance, highlighting a knowledge gap in our understanding of cefiderocol resistance mechanisms. This underscores the need for improved resistance detection strategies in clinical and environmental settings to mitigate the potential dissemination of cefiderocol resistance genes.

## Materials and methods

Soil, wastewater, fish mucus and freshwater samples were collected in Germany, Sweden, Pakistan and France between 2021 and 2022. DNA was extracted using the PowerSoil Pro DNA isolation kit (Qiagen, Hilden, Germany). Detailed methods for each kind of sample can be found in **Supplementary Data**. The quantity and quality of the extracted DNA were assessed using Qubit BR assay along with Nanodrop measurements (Thermofisher, Waltham, USA).

### Metagenomic sequencing and microbial community characterization

Samples were sequenced by the SNP&SEQ Platform at the National Genomics Infrastructure, Uppsala, Sweden. Sequencing libraries were generated using the SMARTer thruPLEX DNA-seq kit (Takara Bio, Shiga, Japan). A total of 47 libraries were pooled across two lanes of an S4 flowcell and were sequenced using the NovaSeq 6000 system. Taxonomic profiling of each metagenome was performed using mOTU v3.1 ^37^. Principal Coordinates Analysis (PCoA) was conducted based on Bray-Curtis dissimilarity between samples. Metagenomic contigs were assembled using NGLess v1.5 and MEGAHIT ^43,44^.

### Functional metagenomic libraries and clone selection

Samples with at least 800 ng of remaining DNA post-sequencing were selected for functional metagenomics detection of cefiderocol resistance genes (see **Supplementary Data** for detailed method). DNA was sheared using the tagmentase enzyme from the Nextera XT kit (Illumina, San Diego, USA) to achieve a target size of 1-3 kb. The tagmented DNA was then amplified and cloned in a pHSG299 expression vector Takara Bio, Shiga, Japan; accession number: M19415). Recombining plasmid was then transformed into competent K12 *E. coli* sensitive to cefiderocol. Resistant clones were then selected on LB agar media supplemented with 100 mM IPTG and 1 mg/L cefiderocol.

### Confirmation of resistance

Confirmation of the cefiderocol-resistant phenotype was achieved by sequencing each clone and assembly of their genomic DNA to identify potential mutations in genes associated with cefiderocol resistance and by transforming fresh competent K12 *E. coli* with the previously extracted plasmid containing the ARG bearing insert (see **Supplementary Data**).

### Phenotypic characterization

Minimum inhibitory concentrations (MICs) were determined in triplicates using unitary UMIC® Cefiderocol tests (Brucker, Billerica, USA) in iron-depleted Mueller Hinton (MH) broth. Disc diffusion assays were also performed to assess susceptibility to a range of beta-lactam antibiotics and beta-lactam/beta-lactamase inhibitor combinations. To further investigate beta-lactamase mediated resistance or the potential effect of the beta-lactamase inhibitor avibactam, cefiderocol + avibactam (4 mg/L) MICs were determined in triplicates using unitary UMIC® Cefiderocol tests (Brucker, Billerica, USA) in iron-depleted MH broth. Additionally, to assess the intrinsic effect of avibactam alone, avibactam MICs were determined in triplicate in MH broth with avibactam concentrations ranging from 0 to 256 mg/L. Finally, a nitrocefin hydrolysis test was also performed.

### Molecular characterization

Initially, the insert amplified via PCR was Sanger sequenced. Taxonomy of the insert sequence was determined using BLASTN against the NCBI ‘nt’ database. ORFs within the insert were identified using PROKKA and BAKTA annotation software ^39,40^. Predicted ARG sequences from the insert were subjected to BLASTN analysis against ARG databases such as ResFinder 4.0 and ResFinderFG v2.0 ^33,38^. BLASTP was also used to identify protein variants within the NCBI protein database ‘nr’. If the resistance mechanism remained unclear, protein structure was predicted using AlphaFold 3 and 3D structure alignments were performed using PyMOL (v3.1.4), both with default parameters ^41^. MatchAlign score divided by the sum of both protein lengths was used to compare alignments. If a functional metagenomic induced cut of the ARG was suspected, the ARG was searched in metagenomic contigs using BLASTN to determine its environment.

### Distribution of cefiderocol-resistant genes

A multi-step bioinformatic approach was employed to determine the environmental distribution and host species of cefiderocol resistance genes identified through functional metagenomics. First, to assess their relative abundance within several environments, metagenomic reads from each sample were mapped against the ARGs using Bowtie2. Next, homologous sequences were identified within the Global Microbial Gene Catalog (GMGC, v1.0; 45)) using a BLAST-like sequence similarity search on GMGC website (https://gmgc.embl.de/; queried on 2025, April 28th). Finally, BLASTN searches against the EnteroBase database were conducted to identify *E. coli* strains harboring these resistance genes.

## Code and data availability

Scripts and software versions used throughout this study are available on the github repository: https://github.com/RemiGSC/Mgf_FDC/ and on the following Zenodo doi: https://doi.org/10.5281/zenodo.15487633. The raw metagenomic and genomic sequencing data generated and analyzed in this study have been deposited in the NCBI Sequence Read Archive (SRA) under the BioProject accession number PRJNA1262354. Accession numbers for whole genome sequencing of cefiderocol resistance clones are: SAMN48516469, SAMN48516470, SAMN48516471, SAMN48516472. The accession number of WGS of the control K12 *E. coli* host transformed with empty pHSG299 is: SAMN48516473.

## Acknowledgments

This study was conducted as part of the “Establishing a Monitoring Baseline for Antibiotic Resistance in Key Environments” (EMBARK) and “Surveillance for Emerging Antimicrobial Resistance through Characterization of the uncharted Environmental Resistome” (SEARCHER) projects, funded by the Swedish Research Council (VR; grants 2019-00299 and 2023-01721), the German Bundesministerium für Bildung und Forschung (F01KI1909A & 01KI2404A), the Agence Nationale de la Recherche (ANR-19-JAMR-0004 and ANR-23-AAMR-0004) under the frame of JPI AMR (EMBARK and SEARCHER; JPIAMR2019-109 and JPIAMR2023-DISTOMOS-016, respectively), the Data-Driven Life Science (DDLS) program supported by the Knut and Alice Wallenberg Foundation (KAW 2020.0239), and the Swedish Foundation for Strategic Research (FFL21-0174). UK & TUB were supported by the Explore-AMR project funded by the German Bundesministerium für Bildung und Forschung (01DO2200). Responsibility for the information and views expressed in the manuscript lies entirely with the authors. We thank Pr André Birgy for his expertise in beta-lactam resistance mechanisms and Dr Luis Pedro Coelho for his advice in the use of GMGC.

## Author contributions

Design of the study was done by R.G. and E.R. Setup of the functional metagenomics was done by R.G., C.D., M.B., I.E. Sampling was done by R.G., A.A., I.D.K., F.T., F.N. and E.R. Metagenomic analysis of each sample was done by R.G., A.A., V.H.J.D., M.W., U.L. and N.G. Functional metagenomics libraries, phenotypic characterization, molecular characterization and confirmation of resistant clones were done by R.G. and M.B. Manuscript writing was done by R.G., M.B. and E.R. Reviewing was done by R.G., M.B., N.G., V.H.J.D., U.L., U.K., R.Z., J.B.P. and E.R. Supervising was done by U.K., T.U.B., S.K.F, R.Z., J.B.P. and E.R.

## Supplementary data

### Supplementary materials and methods

#### DNA isolation

Water and wastewater samples were filtered through 0.22 µm MF-Millipore™ filters (Merck, Darmstadt, Germany) prior to extraction. Filtration volumes varied: 1-6 L for freshwater or saltwater samples, and 0.05-1 L for wastewater input or output. The filters were then shredded and processed according to the kit extraction protocol. In total, 250 mg of material was used per extraction tube for the soil samples. Regarding the fish sample, a European bullhead (*Cottus gobio*) was removed from water of Elbe river in Dresden, Germany as described elsewhere ^58^ and its skin mucus samples taken immediately after the fish was removed from the water using sterile cotton swabs. The collected mucus swab was transferred into a PowerBead tube and stored at −20 °C before DNA extraction. Fishing and animal handling were carried out in accordance with federal legislation and ethics approval based on permits issued by the Saxon State Office for Environment, Agriculture, and Geology (AZ 76/1/9222.22–03/22). Ethical aspects of sampling were conducted following the requirements of Directive 2010/63/EU of the European Parliament and of the Council of 22 September 2010 on the protection of animals used for scientific purposes.

#### Functional metagenomic libraries

DNA was sheared using the tagmentase enzyme from the Nextera XT kit (Illumina, San Diego, USA) following an optimized protocol. To achieve a target size of 1-3 kb, DNA was incubated with a 1/10 dilution of the tagmentase enzyme for 10 seconds at 55°C. The reaction was quickly transferred on ice, and the sheared DNA was purified using QIAquick PCR Purification Kit (Qiagen, Hilden, Germany). The tagmented DNA was amplified using primers designed for adapter ligation, incorporating overlaps necessary for Gibson assembly. DNA fragments within the 1-3 kb range were size selected and purified using QIAquick Gel Extraction Kit (Qiagen, Hilden, Germany).

The expression vector used was pHSG299 (Takara Bio, Shiga, Japan; accession number: M19415). The vector was linearized by PCR using primers designed for adapter ligation, incorporating complementary overlaps required for Gibson cloning.

Gibson assembly was performed using NEBuilder® HiFi DNA Assembly (New England Biolabs, Ipswich, USA). The Gibson assembly reaction mixture was dialyzed against a 0.025 µm MF-Millipore™ membrane (Merck, Darmstadt, Germany). *Escherichia coli* K12 cells were made competent by multiple centrifugation and washes at 4°C in 10% glycerol. A 15 µL aliquot of the Gibson assembly reaction mixture was combined with 35 µL of competent cells, mixed, and then transferred in a 1 mm electroporation cuvette. Cells were electroporated at 1800V and subsequently resuspended in 1 mL Luria-Bertani (LB) media. The cell suspension was incubated for 2 hours at 37°C.

Cells were centrifuged (8000 rpm, 4 min), resuspended in 400 µL of LB medium and 100 µL was used for serial dilutions and plating on LB agar media containing kanamycin (50 mg/L) to estimate library size. The remaining cells were used to inoculate 50 mL LB broth containing 50 mg/L kanamycin for overnight incubation at 30°C. Cells were then centrifuged (8000 rpm, 4 min) and resuspended in 8 mL LB+Glycerol (15%) for storage at -80°C.

#### Clone selection

To select for cefiderocol-resistant clones from the functional metagenomic libraries, *E. coli* carrying libraries was plated on LB agar media supplemented with 100 mM IPTG and 1 mg/L cefiderocol. To ensure the resistance phenotype was attributable to DNA cloned in the expression vector, the cefiderocol resistant colonies were selected as follows: 1) Clones were resuspended in a 20 µL NaCl solution. 2) The bacterial suspension was used to inoculate two separate LB agar plates, one supplemented with kanamycin (50 mg/L) and the other with 100 mM IPTG and 1 mg/L cefiderocol. 3) The remaining bacterial suspension was heated at 95°C and subjected to PCR amplification, targeting the vector insertion site to amplify the inserted DNA fragment. Clones exhibiting growth on both selective media and yielding a positive PCR result were stored at -80°C in LB medium containing 15% glycerol.

#### Confirmation of cefiderocol-resistant phenotype

To further confirm plasmid-mediated cefiderocol resistance, plasmid presumably responsible for the observed cefiderocol MIC were extracted and purified from each resistant clone using the Plasmid Mini Kit (Qiagen, Hilden, Germany). Each purified plasmid was then used to transform fresh competent *E. coli* K12 cells, and a new phenotypic characterization of the transformants was performed.

Additionally, each resistant clone underwent whole genome sequencing (WGS) to discard potential mutations associated with increased cefiderocol MIC. Genomic DNA was extracted using EZ1&2, DNA tissue kit (Qiagen, Hilden, Germany). Libraries for WGS were prepared using Nextera DNA Flex kit and sequenced on a NextSeq (Illumina, San Diego, USA). Reads were quality filtered, and the clone genome was assembled using SPAdes v3.11.1 ^59^. Assembly was annotated using PROKKA v1.14 and potential mutations in genes previously implicated in cefiderocol resistance (detailed in the **Supp Table 2**) were investigated by comparing their sequence to genes found in the *E. coli* K12 transformed with empty pHSG299. Whole genome sequencing data was also analyzed to find potential mutations using Parsnp with default parameters ^60^.

### Supplementary results

#### Confirmation of cefiderocol-resistant phenotype

Fresh K12 *E. coli* transformed with plasmid containing the identified ARG exhibited either cefiderocol MICs that were either equivalent or up to 2 fold higher than those of the original cefiderocol resistant clone. than the one observed for the original resistant clone. Antibiotic disc diffusion assays also confirmed that both clones displayed the same resistance phenotype.

Subsequent genomic DNA sequencing of each resistant clone revealed no mutations in genes previously associated with cefiderocol resistance (**Supp table 3**). Instead, random mutations were found in specific clones within intergenic regions and in genes encoding rhsC (a transposase), GTPase Der and a primosomal replication protein N.

---

**Supp Table 1:** Alignment of the ARGs identified by functional metagenomics using BLASTN against Enterobase database. Only results with pident > 95% and qcovhsp >95% are shown.

| ARG | *E. coli* strain | pident | qcovhsp | mismatch | gapopen | eval | bitscore | Serotype | ST | fimH | Phylogroup |
| --- | --- | --- | --- | --- | --- | --- | --- | --- | --- | --- | --- |
| GER1FDC1 | ESC_KA9999AA_AS | 97,9 | 100 | 19 | 0 | 0.0 | 1569 | O26:H32 | ST10/- | fimH24 | A |
| GER1FDC1 | ESC_IA4897AA_AS | 97,9 | 100 | 19 | 0 | 0.0 | 1569 | O128:H12 | ST472/- | fimH457 | A |
| GER1FDC1 | ESC_IA4024AA_AS | 97,9 | 100 | 19 | 0 | 0.0 | 1569 | O129:H30 | ST176/- | fimH23 | A |
| GER1FDC1 | ESC_NA7870AA_AS | 97,9 | 100 | 19 | 0 | 0.0 | 1569 | Unknown:H4 | ST88/74 | fimH39 | C |
| GER1FDC1 | ESC_IA4025AA_AS | 97,9 | 100 | 19 | 0 | 0.0 | 1569 | O129:H30 | ST176/- | fimH23 | A |
| GER1FDC1 | ESC_IA6016AA_AS | 97,9 | 100 | 19 | 0 | 0.0 | 1569 | O129:H30 | ST176/- | fimH23 | A |
| GER1FDC1 | ESC_LA1083AA_AS | 97,9 | 100 | 19 | 0 | 0.0 | 1569 | O129:H30 | ST176/- | fimH23 | A |
| GER1FDC1 | ESC_VA2039AA_AS | 97,8 | 100 | 20 | 0 | 0.0 | 1564 | O179:H9 | ST410/471 | fimH24 | C |
| GER1FDC1 | ESC_RA9125AA_AS | 97,8 | 100 | 20 | 0 | 0.0 | 1564 | O179:H9 | ST-/471 | fimH24 | C |

ARG: antibiotic resistance gene; pident: percentage of identical matches; qcovhsp: query coverage per High-Scoring Pair (HSP); ST: sequence type; FimH: Type 1 fimbriae adhesin.

**Supp Table 2:**
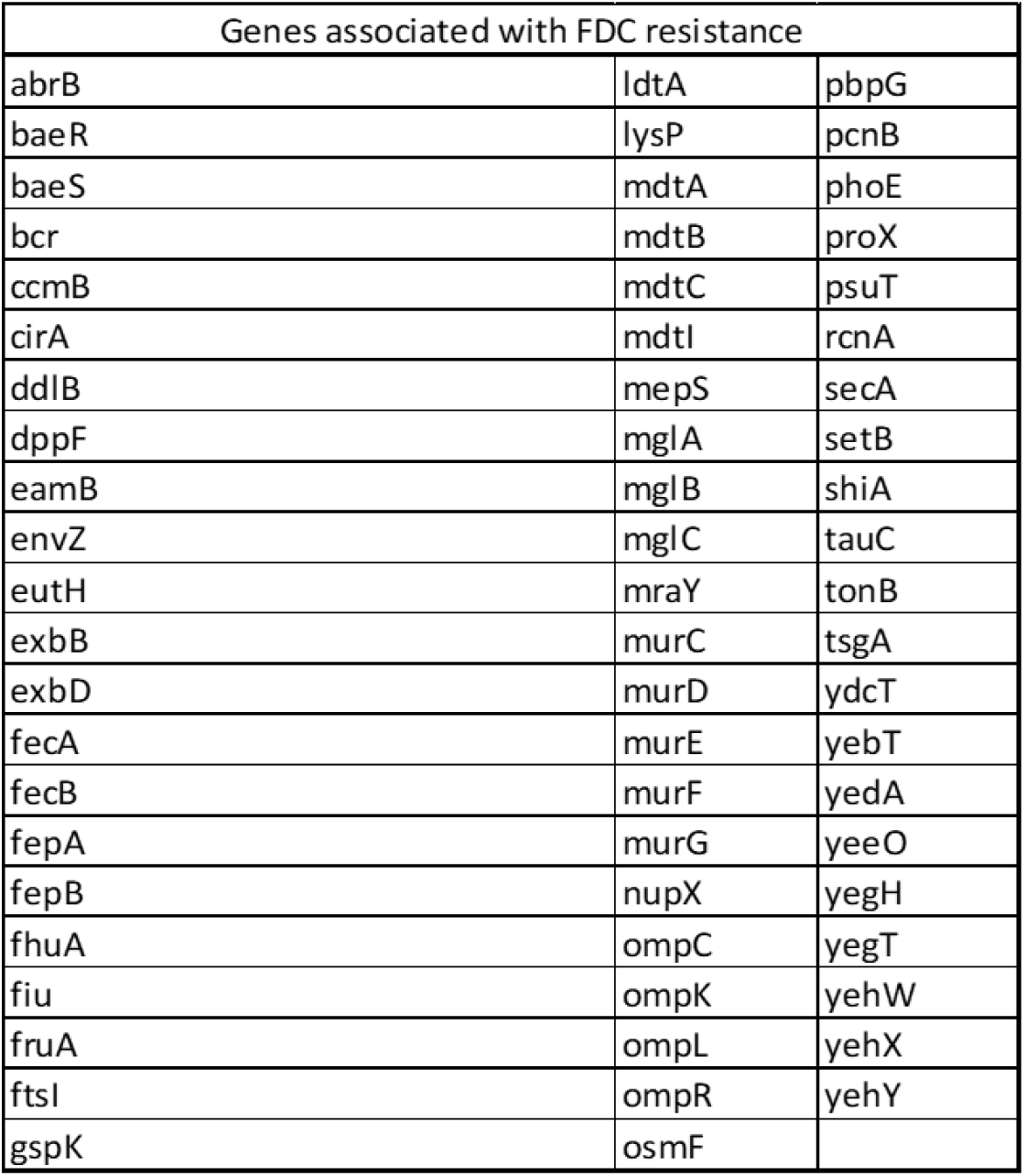
Genes involved in the check of mutation or modifications on cefiderocol resistance associated genes.

**Supp table 3:**
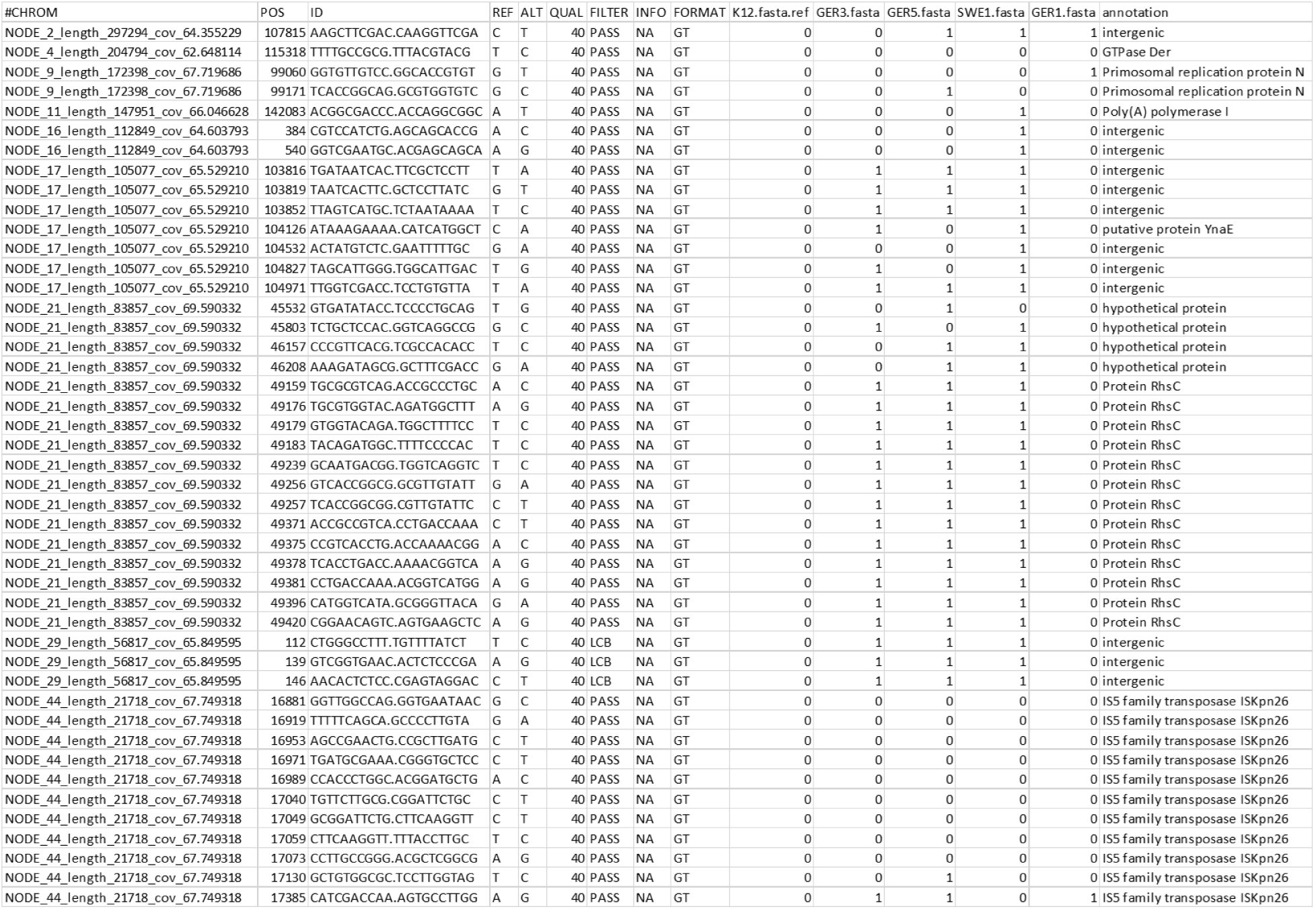
Mutations identified between *E. coli* K12 used for functional metagenomics and each cefiderocol resistant clone.

**Supp Figure 1:**
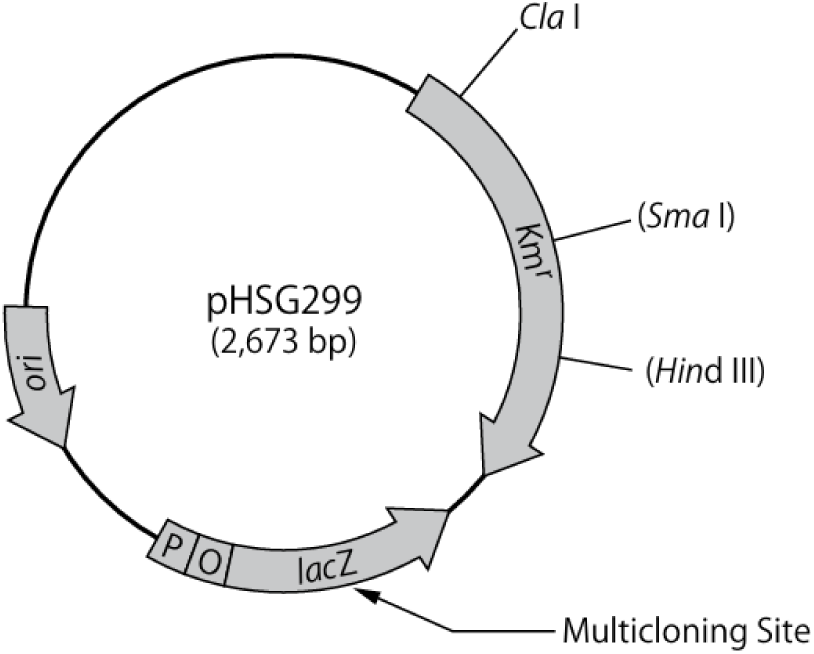
cartography of pHSG299 plasmid. Figure comes from the manual which can be found on TAKARA bio website (https://www.takarabio.com/documents/).

**Supp Figure 2:**
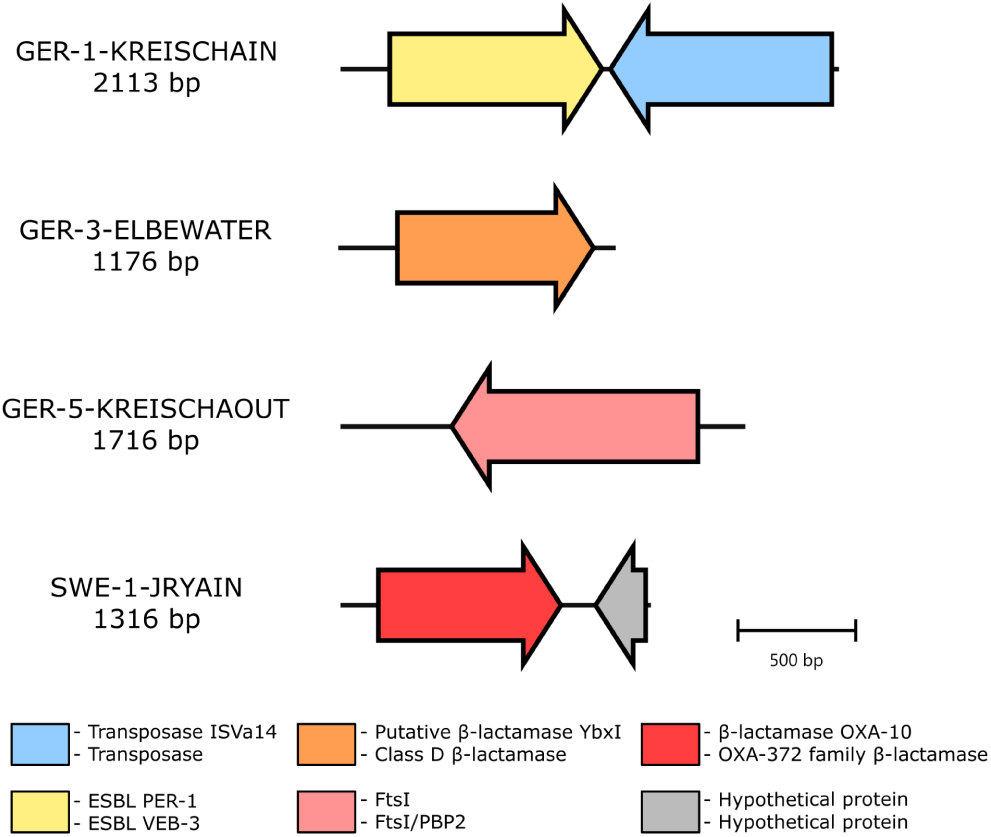
cartography of insert sequences associated with elevated cefiderocol MICs and identified using functional metagenomics. ORFs and gene annotations were obtained using PROKKA v1.14 (up) and Bakta web server v1.11.0 (down). bp: base paire.

**Supp Figure 3:**
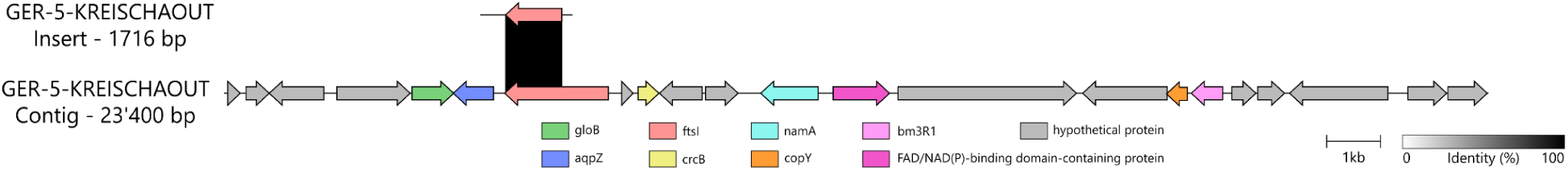
cartography of the insert sequence identified in the GER-5-KREISCHAOUT sample and of the contig from metagenomic sequencing reads assembly of the same sample where the gene could also be found. Gene annotations were obtained using Bakta web server v1.11.0.

